# Cryo-electron tomography reveals coupled flavivirus replication, budding and maturation

**DOI:** 10.1101/2024.10.13.618056

**Authors:** Selma Dahmane, Erin Schexnaydre, Jianguo Zhang, Ebba Rosendal, Nunya Chotiwan, Bina Kumari Singh, Wai-Lok Yau, Richard Lundmark, Benjamin Barad, Danielle A. Grotjahn, Susanne Liese, Andreas Carlson, Anna K. Överby, Lars-Anders Carlson

## Abstract

Flaviviruses replicate their genomes in replication organelles (ROs) formed as bud-like invaginations on the endoplasmic reticulum (ER) membrane, which also functions as the site for virion assembly. While this localization is well established, it is not known to what extent viral membrane remodeling, genome replication, virion assembly, and maturation are coordinated. Here, we imaged tick-borne flavivirus replication in human cells using cryo-electron tomography. We find that the RO membrane bud is shaped by a combination of a curvature-establishing coat and the pressure from intraluminal template RNA. A protein complex at the RO base extends to an adjacent membrane, where immature virions bud. Naturally occurring furin site variants determine whether virions mature in the immediate vicinity of ROs. We further visualize replication in mouse brain tissue by cryo-electron tomography. Taken together, these findings reveal a close spatial coupling of flavivirus genome replication, budding, and maturation.

## Introduction

*Orthoflaviviruses* (henceforth flaviviruses) are a large genus of arthropod-borne, positive-sense RNA viruses within the *Flaviviridae* family. The mosquito-borne Dengue virus alone is estimated to yearly cause hundreds of millions of human infections, some progressing to the severe condition known as dengue shock syndrome^1^. Human infections with tick-borne flaviviruses are less frequent, but can have severe outcomes. Tick-borne encephalitis virus (TBEV) is the namesake virus of the “TBEV serocomplex” which includes other tick-borne flaviviruses such as Powassan virus and the low-pathogenic Langat virus (LGTV). Pathogenic tick-borne flaviviruses have a strong neurotropism in mammals, and can cause encephalitis with debilitating or deadly outcome in humans^2^.

After entering the cell through endocytosis, the flavivirus genome is translated as a single, transmembrane polyprotein, which is subsequently cleaved by host and viral proteases into ten individual proteins. Seven of these are the non-structural (NS) proteins, which serve to replicate the viral genome. Of the NS proteins, NS3 and NS5 are cytoplasmic enzymes that serve as protease and helicase (NS3), and RNA-dependent RNA polymerase and methyl transferase (NS5). The remaining NS proteins include the endoplasmic reticulum (ER) lumen-resident peripheral membrane protein NS1, and the integral membrane proteins NS2A, NS2B, NS4A and NS4B. Viral genome replication takes place on a transformed, dilated ER containing multiple bud-like membrane invaginations^3,4^. These invaginations, referred to as replication organelles (ROs), are the site of viral RNA replication^5-9^. RO-like membrane rearrangements can be formed by a subset of NS proteins even in the absence of viral RNA replication^5,10,11^, but require interactions with host ER proteins^5-9,12,13^. Electron microscopy of resin-embedded, infected cells has shown that the RO is a 80-90 nm, near-spherical bud with a ∼10 nm opening towards the cytoplasm^14-17^. However, due to the destruction of protein structure by resin embedding, the organization of proteins and RNA in the RO is still unknown. Virion assembly also takes place at the ER, when a cytoplasmic complex of viral RNA and C protein interacts with the transmembrane envelope proteins prM and E, followed by budding into the ER lumen. NS2A has been suggested as a key viral protein coupling replication and assembly^18-20^, and resin-embedding electron microscopy has visualized putative virions in the immediate vicinity of ROs^14,16,21^. Newly formed, immature virions have a spiky surface covered with extended prM-E trimers^22-24^. The cleavage of prM by the host-cell protease furin, which is thought to occur in the trans-Golgi, leads to a structurally rearranged, infectious, mature virion with smoother appearance^25,26^. If virion assembly and maturation are directly linked to ROs is currently unknown.

To shed light on the interactions between flavivirus replication, assembly and maturation, we performed *in situ* cryo-electron tomography^27-31^ on human cells and mouse brain tissue infected with LGTV, and a novel, chimeric LGTV carrying TBEV structural proteins. The data suggest a mechanism for RO membrane remodeling, the presence of a protein complex tethering the RO membrane to an apposed ER membrane, and a close proximity of virion assembly and maturation.

## Results

### Cryo-electron tomography reveals two states of replication organelles in Langat virus-infected cells

To explore the macromolecular architecture of flavivirus ROs, we grew human A549 cells on EM grids and infected them with LGTV. Cells were plunge-frozen at 24 h post infection (h.p.i) and subjected to focused-ion-beam milling, after which lamellas containing the infected cytoplasm were imaged using cryo-electron tomography (cryo-ET) (Table 1). The tomograms revealed a dilated ER inclusive of clustered ROs (Fig. 1 and Supplementary video 1). ROs were clearly identified as near-spherical membrane invaginations into the ER lumen, of a kind not present in uninfected cells (Fig. S1A-B). The vicinity of the remodeled ER contained *bona fide* ribosomes as well as mitochondria immediately apposed to the ER membrane (Fig. 1A-D). ROs frequently appeared in clusters within the lumen of dilated ER, as in Fig. 1A-D in which a single, dilated ER cisterna contained >10 ROs within the field of view (bearing in mind that the RO cluster probably extended beyond the depth of the lamella). The same ER cisterna additionally contained a virus particle with the characteristic spiky appearance of immature flaviviruses (Fig. 1A-B, orange arrow, and Fig. 1D). The majority of ROs contained filamentous densities, presumably the replicating double-stranded form of the viral RNA, within their lumen (Fig. 1A-D). On the other hand, several ROs were devoid of internal filamentous structures (Fig. 1A-C, white arrows). These two types of ROs will henceforth be denoted as ‘filled’ and ‘empty’, respectively (Fig. 1E). In 23 tomograms, 83±14% of ROs were filled (Fig. 1F). The empty ROs were significantly smaller with an average diameter of 46±8 nm (N=25), compared to the filled ROs at 85±5 nm (N=63) (Fig. 1G). In summary, we established a workflow to image flavivirus replication by cryo-ET, revealing that ROs exist in two forms: with and without luminal filamentous densities.

**Table 1.** Number of tilt series recorded and tomograms analyzed. “Number of tilt series” refers to the total number of tilt series recorded on a given sample. All of these tilt series were used to reconstruct tomograms, and the “number of tomograms with events” refers to the number of tomograms with virus replication-related events.

| sample | number of tilt series | number of tomograms with events |
| --- | --- | --- |
| LGTV WT | 51 | 23 |
| rLGTV<sup>T:prME</sup> R86 | 12 | 7 |
| rLGTV<sup>T:prME</sup> Q86 | 18 | 7 |
| LGTV in *ex vivo* choroid plexus | 45 | 8 |

**Figure 1:**
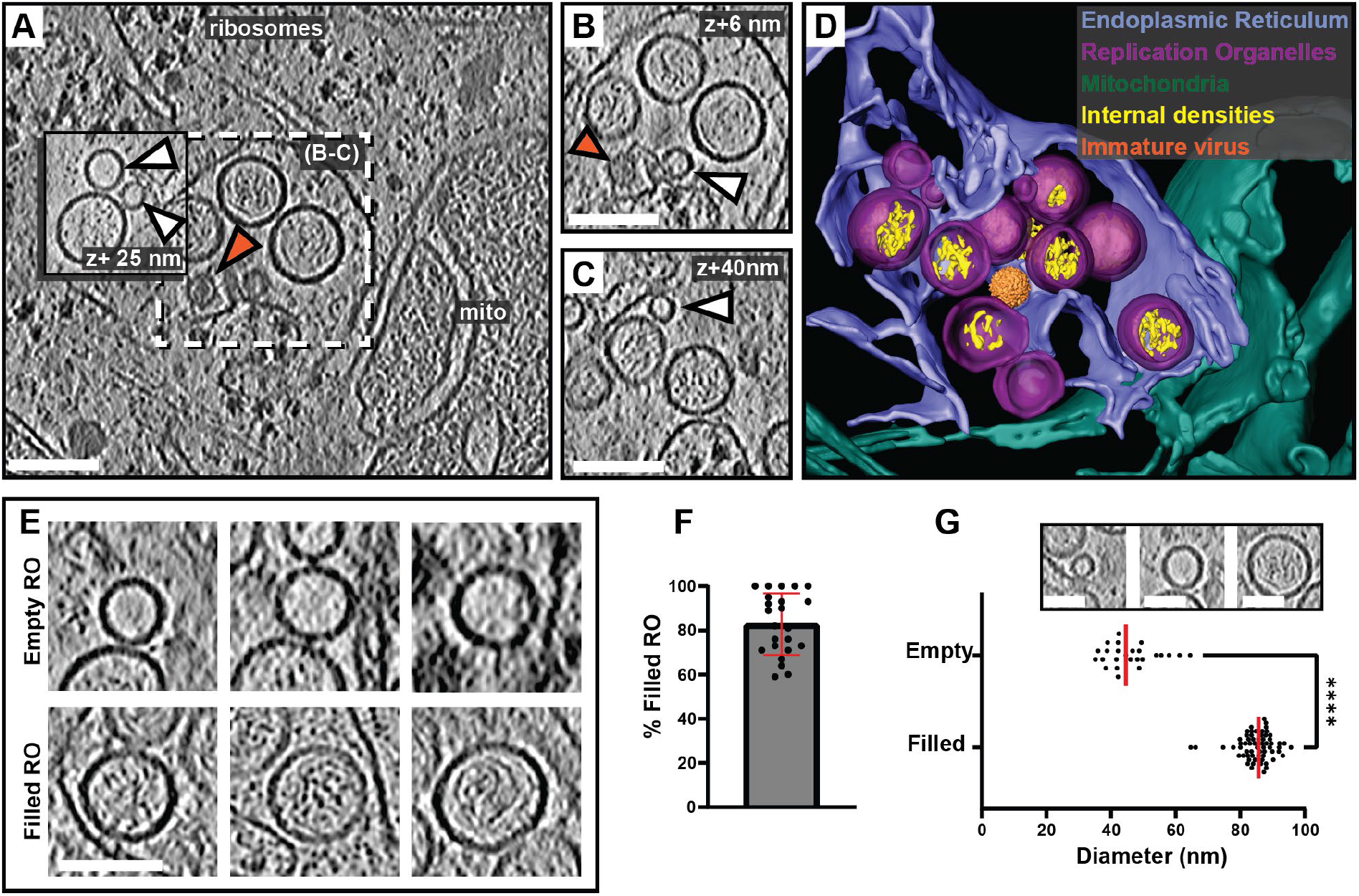
*In situ* cryo-ET uncovers two states of Langat virus replication organelles. (A) Slice through a tomogram of FIB milled LGTV-infected cell showing viral ROs enclosed within the ER with an immature virion (orange arrow). (B-C) Close up views of the outlined framed region in (A) at their respective Z heights in the tomogram. (A-C) White arrows indicate empty ROs. (D) Segmentation of the tomogram in (A), with color schemes defined for each structure. The immature virion is represented by a subtomogram average. (E) Representative examples of empty and filled viral RO observed in cryo-tomograms of milled LGTV-infected cells. (F) Percentage of filled ROs observed in 23 tomograms of LGTV-infected cells. (G) Size distribution of empty (n=25) and filled (n=63) RO observed in tomograms. The inset represents the different sizes of viral spherules observed in the tomograms. (F-G) Red lines represent the average, each dot corresponds to one analyzed tomogram (see also Table <u>1</u>). Statistical significance by unpaired two-tailed Student’s *t* test: \*\*\*\**p* < 0.0001. Scale bars 100 nm.

### A combination of membrane coat and RNA-induced pressure determines RO morphology

ROs of other viruses depend on the pressure from intraluminal double-stranded RNA (dsRNA) to inflate and stabilize the curved RO membrane^32^. The presence of empty LGTV ROs speaks against this mechanism for flaviviruses, and we thus reasoned that a coat might confer a spontaneous curvature to the RO membrane. To investigate whether a curvature-inducing coat is present on ROs, we assessed if RO membrane thickness is in line with a layer of surface-bound protein layer. Indeed, by visual inspection of tomograms, the membranes of both empty and filled ROs appeared thicker than the surrounding ER membrane (Fig. 2A). No distinct, repeating macromolecules were visible on the RO membranes. While this does not exclude the presence of a protein coat composed of smaller or membrane-integral proteins, it does make its characterization by subtomogram averaging less likely to be successful. Instead, we extended our previously developed surface morphometrics toolbox to allow for local estimation of membrane thickness^33^. This software allowes calculation of the average membrane thickness within a user-selected area, based on average density profiles normal to the membrane. Both for single ROs and their surrounding ER membrane, this yielded reliably interpretable density profiles (Fig. 2B-C). Color coding membranes by thickness indicated that RO membranes are consistently thicker than the surrounding ER membrane (Fig. 2D). Indeed, in four tomograms we measured a significant difference in membrane thickness with the ER membrane being 3.4 ± 0.2 nm (N= 4), and RO membranes 4.0 ± 0.2 nm (N =132) (Fig. 2E). On the other hand, the RO membrane thickness appeared largely independent of RO size, as estimated from their radii of curvature (Fig. 2F).

**Figure 2:**
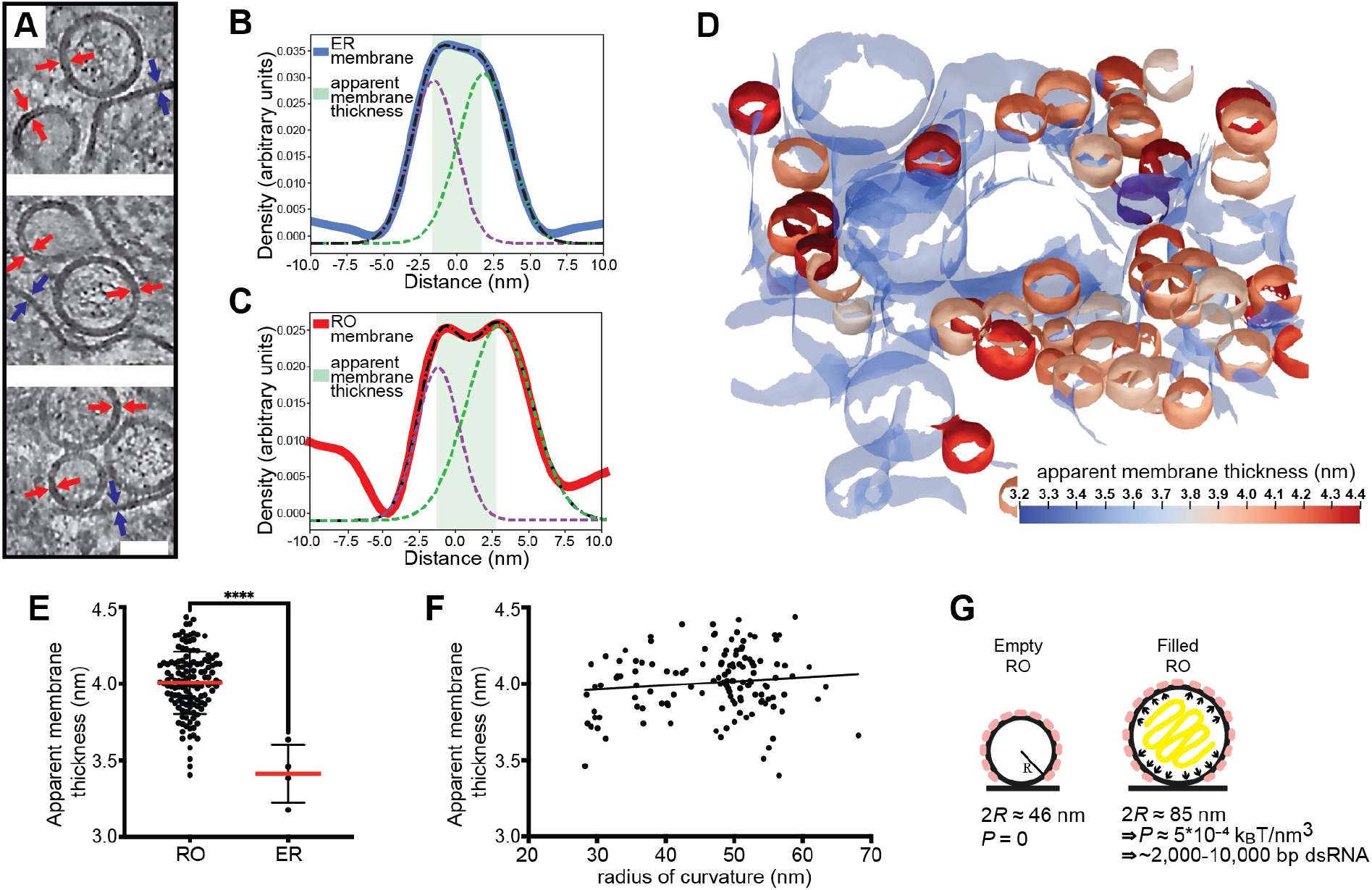
The influence of a membrane coat and viral RNA in shaping the replication organelles. (A) Slices through tomograms of empty and filled ROs in LGTV-infected cells. Blue and red arrows indicate the thickness of the ER and RO membranes, respectively. Scale bar, 50 nm. (B-C) Membrane thickness estimation by dual Gaussian fitting to radial density plots through a representative ER membrane (B) and RO membrane (C). Solid lines, membrane density profile; dashed lines, fitted composite and dual Gaussian; shaded area, estimated thickness. (D) ER and ROs membranes from a representative tomogram of an LGTV-infected cell, color coded by apparent membrane thickness. ER membrane is partially transparent. (E) Apparent membrane thickness quantification in four tomograms of LGTV-infected cells, comparing individual ROs (n=132) and the surrounding ER (n=4). Red lines, average. Statistical significance by unpaired two-tailed Student’s t test: ****p < 0.0001. (F) Relationship between radius of curvature and apparent membrane thickness for individual ROs (n=132 from six tomograms). (G) Model of the mechanisms determining viral RO size. Two RO states exist in infected cells: empty ROs with a baseline size set independent of luminal RNA, and filled ROs, whose larger size is due to intraluminal pressure from ∼2,000-10,000 bp dsRNA.

The observation that flavivirus ROs can form without detectable luminal dsRNA, and the consistent presence of a membrane coat, distinguish them from alphavirus ROs that have a near-identical membrane shape. Based on cryo-ET data, we recently published a mathematical model of alphavirus RO membrane budding, which showed that the pressure from intraluminal dsRNA, together with constraint of the membrane neck, is sufficient for the creation of the RO membrane bud^32^. We next adapted this mathematical model to explain flavivirus RO membrane remodeling. We assume that the membrane coat generates a spontaneous curvature *H*_0_ of the RO membrane. Such a spontaneous curvature can be generated by the protein structure as well as by crowding^34^. Furthermore, we consider that in ROs that enclose dsRNA, the RNA exerts a pressure *P* on the membrane. The total energy *E* of the RO membrane is then composed of an integral over the membrane surface *A* and the contribution of the pressure, which scales with the volume *V*,

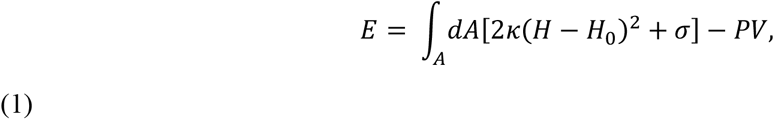

where the first term describes the bending energy according to the Helfrich model, with *κ* the bending stiffness and *H* the mean curvature^35^. The second term in Eq. 1 contains the membrane tension *σ*. Motivated by the experimentally observed shapes, we describe the ROs as spheres with a radius *R*, as schematically depicted in Fig. 2G, simplifying Eq. (1) to

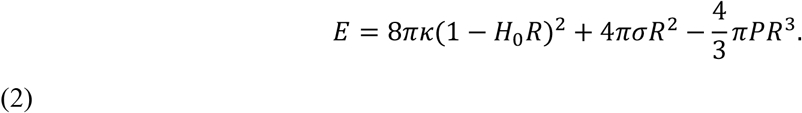

However, there are two unknown factors in Equation (2), the spontaneous curvature *H*_0_ and the pressure P in the RO. Based on our observations above, the membrane coat can be assumed to be comparable for empty and filled ROs. Thus, we can take advantage of the imaging of empty ROs to obtain *H*_0_ at vanishing pressure, *P* = 0. Minimizing Eq. (2) with respect to *R*, we obtain

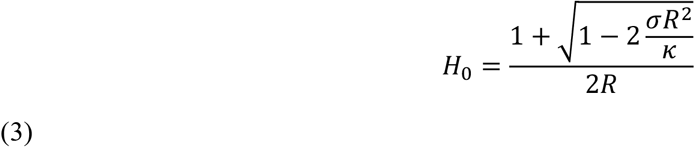

From the experiments we have obtained an average diameter of the empty ROs to be 2*R* = 46 *nm*. By using previously estimated parameters^32^ for the membrane properties, i.e., *σ* = 10^−5^*N/m* and *κ* = 10*k*_*B*_*T*, we predict the spontaneous curvature to be *H*_0_ = 0.04*nm*^−1^, which corresponds to a radius of curvature 1*/H*_0_ = 25*nm*. Next, we want to predict the influence of the pressure generated by the RNA on the spherule size. Since the RNA does not affect the spontaneous curvature *H*_0_, we keep the above prediction and minimize Eq. 2 with respect to *R*, giving

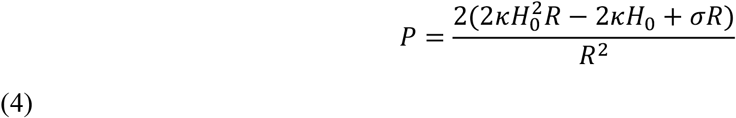

Now including the predicted *H*_0_ = 0.04*nm*^−1^ with the measured average diameter of RNA-filled ROs 2*R* = 85 *nm*, we obtain the pressure *P* = 5 · 10^−4^ *k*_*B*_*Tnm*^−3^. To interpret this value, we compare it with our previous study of alphavirus ROs, which shows that a dsRNA with a length of 2,000-10,000 base pairs generates an internal pressure of 10^−4^ − 10^−3^ *k*_*B*_*Tnm*^−3^ (see Materials and Methods, section Estimating RO intraluminal pressure).

Taken together, flavivirus ROs have a consistently thicker membrane than the surrounding ER, consistent with the presence of small, curvature-stabilizing proteins that set a baseline RO size in the absence of luminal RNA. The size increase from empty to filled ROs is consistent with a single copy of the genome in dsRNA form.

### Virions form and undergo maturation in the immediate vicinity of replication organelles

In our cryo-electron tomograms we consistently observed virions in the vicinity of ROs, underscoring the strong association between replication and virion assembly (Fig. 3A-C). We next wished to use the structure preservation in cryo-ET to study the spatial relation between virion budding and maturation. In the tomograms, virus particles at different stages of maturation were distinguishable: spiky particles corresponding to immature virions as well as smooth, mature virions (Fig. 3A-C). Subtomogram averaging on a small number of virions confirmed the distinct morphology of the immature and mature virions (Fig. 3D-E), and a good match between the *in situ* averages and structures of purified flaviviruses (Fig. S3). The tomograms also included examples of what seemed to be nearly or recently completed virion budding (Fig. 3A-B). In such events, immature virus particles could be observed right at the membrane, across from ROs (Fig. 3A, orange arrow). Both immature and mature virions were consistently found close to ROs, in separate but intertwined membrane compartments (Fig. 3A-C). While immature and mature particles were not observed in the same membrane compartment, seemingly discrete compartments may have been connected beyond the limited thickness of the lamella. To quantitate the relation of immature and mature particles to ROs, we measured the center-center distance of virions to ROs in 5 tomograms. Immature virions were 95±56 nm (N=37), and mature virions 147±41 nm (N=24) from the closest RO (Fig. 3F). The small but significant distance difference (p=0.0003, unpaired t test), together with the observation that immature and mature virions are present in separate compartments, indicates that virion maturation is coupled to a slight spatial separation from ROs but does not necessitate longer-range trafficking. To further investigate the spatial relationship between genome replication, virion assembly and the Golgi apparatus, we performed immunofluorescence light microscopy on fixed, LGTV-infected cells. Furin exhibited co-localization with the Golgi marker GM130 in both uninfected (Fig. S4A-D) and LGTV-infected cells (Fig. 3G-J). Meanwhile, the dsRNA signal was detected in close proximity to and partially overlapping with clusters enriched in furin and GM130 (Fig. 3J). These results corroborate the cryo-ET, reinforcing the close proximity between viral RNA replication, LGTV assembly and maturation within the infected cell. In summary, both virion assembly and maturation can occur in the immediate proximity of ROs, in distinct but intertwined compartments.

**Figure 3:**
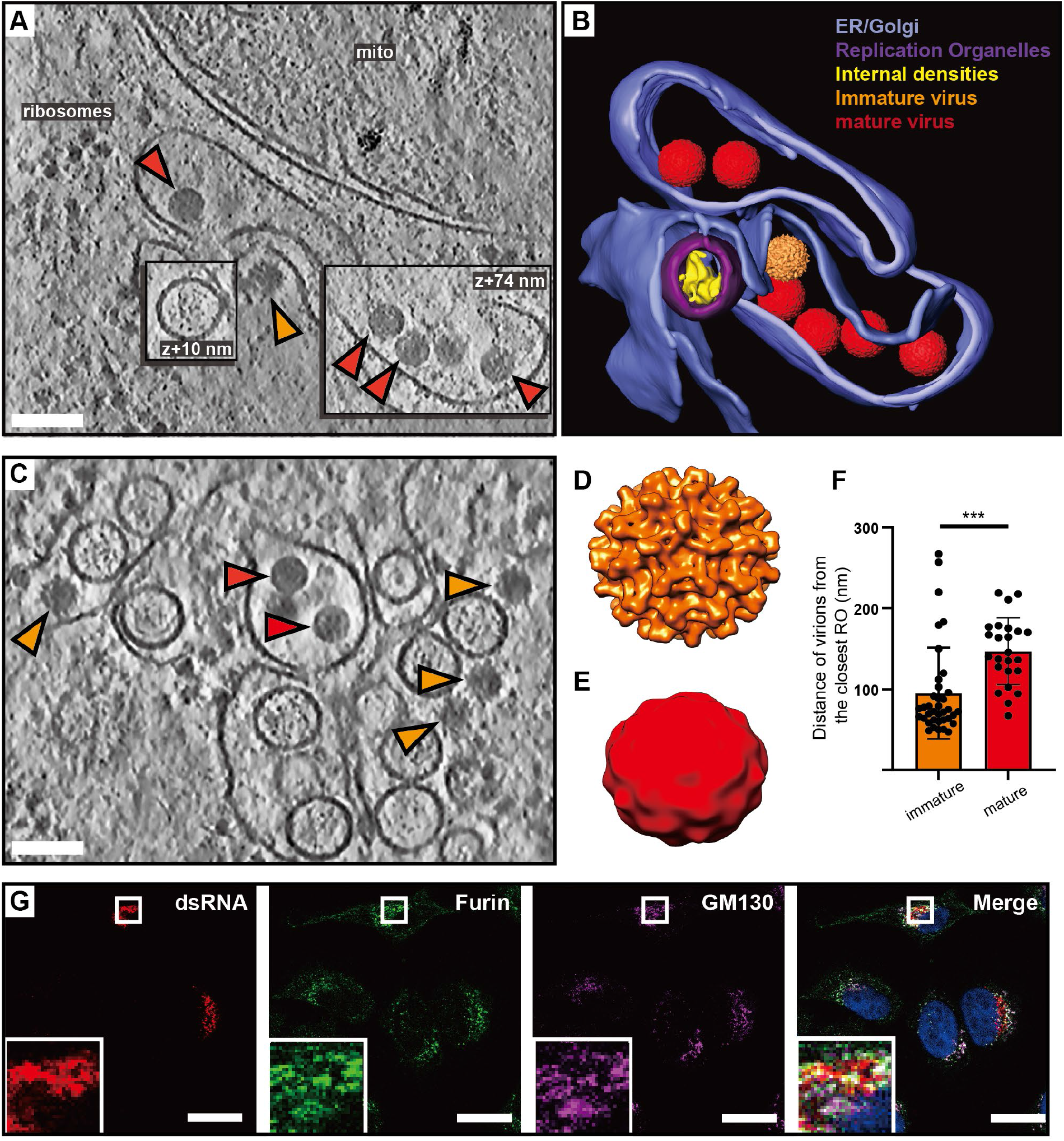
Virions form and mature in the immediate vicinity of replication organelles. (A) Slice through tomogram of LGTV-infected cell showing an immature virion budding across from a RO (orange arrow), and mature virions (red arrows) in adjacent membranes. (B) Segmentation of the tomogram in (A), with color labels defined for each structure. (C) Slice through tomogram of LGTV-infected showing mature (red arrows) and immature (orange arrows) LGTV virions observed near the viral RO. (D-E) Subtomogram averages of immature (D) and mature (E) LGTV from cellular tomograms. (F) The distance from immature (n=37) and mature (n=24) virions to the closest RO, from five tomograms. Statistical significance by unpaired two-tailed Student’s *t* test: \*\*\*\**p* < 0.005. (G) Representative immunofluorescence micrograph of dsRNA, furin and Golgi marker GM130 in LGTV-infected cells at 24 h p.i. Rightmost panel: merge including DAPI-staining of nuclei (blue). Scale bars 100 nm (A,C), 5 µm (G).

### The TBEV furin site variants R86 and Q86 differ in replication organelle-proximal virion maturation

In purified virions, the transition from immature to smooth conformation can be brought about by pH change even in the absence of furin cleavage. Thus, we wanted to test if the observed RO-proximal virion maturation was dependent on the furin site in prM. To do so, we took advantage of our recently characterized chimeric LGTV, which carries the structural proteins prM and E(ectodomain) from TBEV (Fig. 4A). This virus, rLGTV^T:prME^, is genetically stable and has a low pathogenicity similar to that of wildtype LGTV^36^. The rLGTV^T:prME^ prM and E come from the human TBEV isolate 93/783 of the European subtype, whose prM has an unusual arginine (R) in position 86 where most other strains have a glutamine (Q) (Fig. 4A). This residue is located at the P8 position of the furin cleavage site in prM, and previous studies of flaviviruses has shown that modifications here might influence prM cleavage and virus export^37^. We thus reasoned that R86 may affect furin cleavage efficiency, and provide a naturally occurring tool to study maturation. Hence, we also produced a version of rLGTV^T:prME^ with the more common glutamine 86. These chimeric viruses, henceforth referred to as R86 and Q86, only differ in this amino acid residue. Both viruses replicated with similar kinetics in A549 cells (Fig. S5A), but the R86 virus had a slightly higher lethality in immunocompromised *Ips1*^-/-^ (IFN-β promoter stimulator 1) mice (5 of 5 mice died with R86, 7 of 10 died with Q86, Fig. 4B). No difference in neurovirulence was detected (Fig. 4C). We next wanted to investigate whether the pathogenicity of R86 as compared to Q86 correlated with different prM cleavage kinetics. In a cleavage assay with a peptide corresponding to residues 81-94 of prM, the R86 sequence was cleaved faster by furin than Q86 (Fig. 4D). We noted that the R86 sequence generates a putative second, minimal recognition site (KR) for other proprotein convertases such as PC1/3 and PC2^38^. In a peptide cleavage assay, the R86 sequence was also cleaved faster than Q86 by PC1/3. However, the cleavage was still completely dependent on the furin recognition site (RTRR), i.e. the putative second PC1/3 cleavage site K85-R86 was not sufficient for cleavage by PC1/3 (Fig. 4D). Next, we looked for the presence of unprocessed prM protein in cell supernatant, which would indicate release of immature virus particles. For both viruses, the bulk of cell-bound M was in the form of uncleaved prM, whereas most M in the supernatant was cleaved (Fig. 4E). Interestingly, the R86 chimeric virus had a significantly lower percentage of uncleaved prM in supernatant at 48 h post infection (Fig. 4F), suggesting that R86 confers a more efficient particle maturation. The biochemical and cell assays thus converge on the interpretation that the R86 sequence variant confers faster furin cleavage.

**Figure 4:**
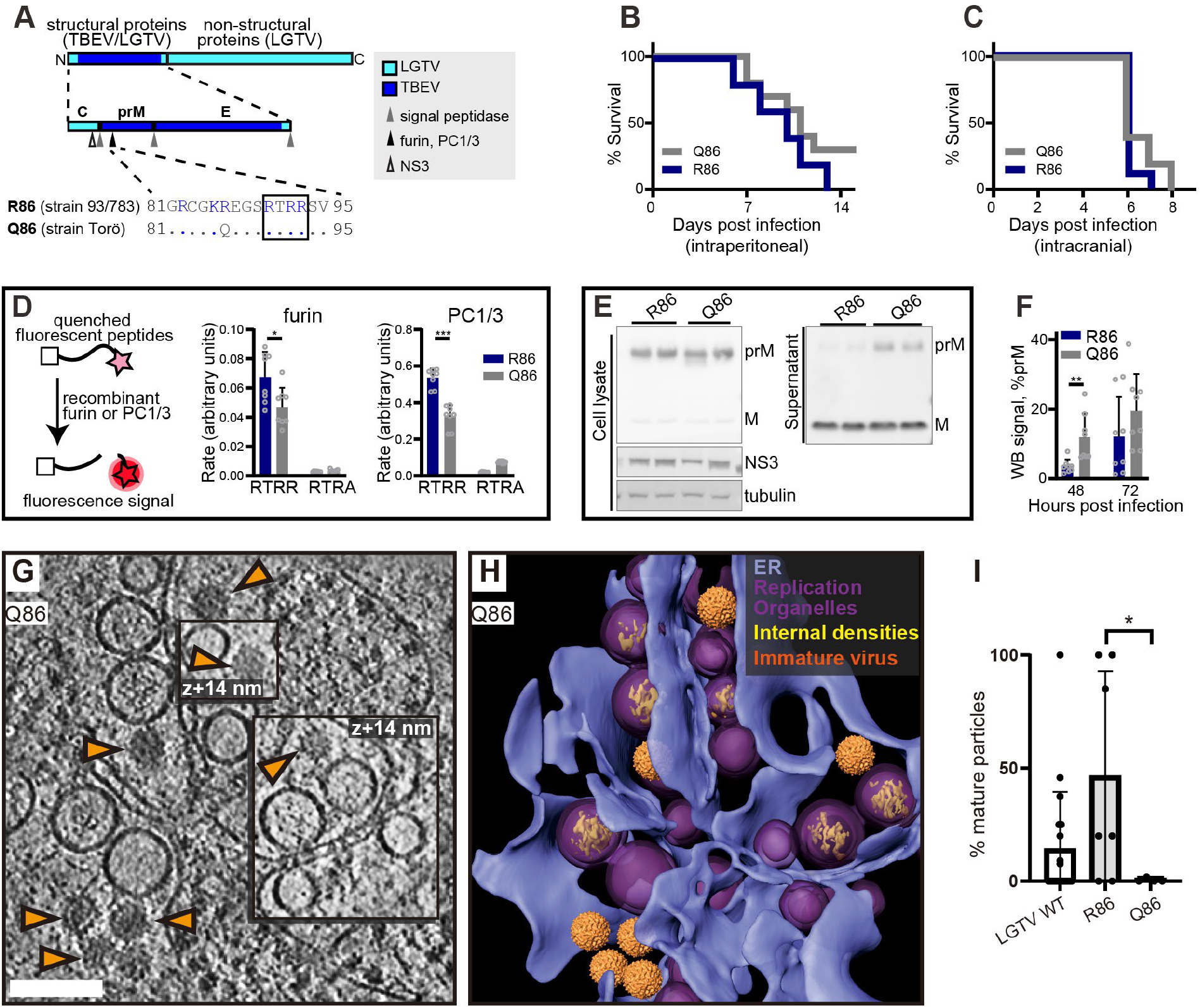
The TBEV furin site variants R86 and Q86 differ in cleavage efficiency and replication organelle-proximal virion maturation. (A) The polyprotein of a chimeric LGTV with prM and ecto-E from TBEV strain 93/783, as color coded. Recognition sites for viral and cellular proteases are shown within the structural protein region, and the furin site sequences from TBEV strains 93/783 and Torö are shown highlighting the difference at position 86 (., identical sequence). (B) Percentage survival of *Ips1*^-/-^ mice infected intraperitoneally with 10^4^ FFU with chimeric LGTV R86 (n=5) or Q86 (n=10). (C) As (B), for but *Ips1*^-/-^ mice infected intracranially with 10^2^ FFU (n=9, R86 and n=10, Q86). (D) Enzymatic cleavage using recombinant furin or PC1/3 with peptides corresponding to furin site sequences in (A) (RTRR), or peptides with impaired furin sites (RTRA). Data from four independent experiments performed in duplicates are shown, with mean values and standard deviation. (E) prM and M protein levels in cell lysates and supernatant 48 h.p.i. visualized by immunoblotting. Viral NS3 and cellular tubulin included as infection and loading control. Representative blots are shown. (F) Ratio of prM/M intensity quantified in supernatants at 48 and 72 h.p.i. Data from four independent experiments performed in duplicates are shown, with mean values and SD. (G) Slice through tomogram of chimeric LGTV Q86-infected cell showing a predominance of immature virions (orange arrows) within the cytoplasm. Scale bar 100 nm. (H) segmentation of the tomogram in (A), with color labels defined for each structure. (I) Percentage of mature virions in the tomograms of LGTV WT (n=19), chimeric LGTV R86 (n=6), LGTV Q86-infected cells (n=4). (D,F,I) Statistical significance by unpaired two-tailed Student’s *t* test: *p<0.05, **p<0.01, ***p < 0.001.

Having characterized the different rates of furin cleavage, we then returned to the question of individual virion conformation inside infected cells. We infected cells with R86 and Q86 chimeric viruses and recorded cryo-electron tomograms of the infected cytoplasm as for wildtype LGTV. Both for R86 and Q86, the tomograms showed a similar overall appearance as for wildtype LGTV, including an abundance of filled and empty replication organelles as well as new virions inside dilated ER compartments (Fig. 4G-H, Fig. S5B). In the R86 tomograms, both immature and mature virions were seen, whereas immature virions appeared to predominate in the Q86 tomograms. We calculated the percentage mature virions in a set of tomograms for wildtype LGTV and the chimeric viruses. Whereas the number of virions in individual tomograms was sometimes small, the average fractions mature particles over several tomograms showed a clear trend (Fig. 4I). For Q86, 2.5±5.9% of virions were mature (N=7 tomograms), whereas 46±46% of R86 virions were mature (N=7 tomograms), a significantly higher percentage (p=0.02, unpaired t test). Wildtype LGTV was intermediate to the two recombinant viruses, with 14±25% mature virions. Taken together, a single residue in the distal part of the TBEV furin site affects prM cleavage rates by furin and PC1/3, mean survival in immunocompromised mice, and particle maturation in areas near ROs.

### A protein complex connects the replication organelle to an apposed ER membrane

A recurring feature in the cryo-electron tomograms was the close proximity of a second ER membrane to the ER membrane containing the ROs (Figs. 1-2). We noticed that this was consistently mediated by a protein complex present at the membrane neck of the replication organelle, connecting this membrane to the adjacent, second ER membrane (Fig. 5A-D). While limited occurrences of these complexes hindered structural analysis, they consistently appeared at the necks of both filled and empty ROs (Fig. 5A-D). From volume estimates of individual complexes, we estimate their molecular masses to 500±151 kDa (Fig. 5E). Interestingly, we observed similar-looking complexes connecting the replication organelle to the site of immature particle budding in the neighboring ER cisterna (Fig. 5F-H). This observation suggests that this protein complex might play a role in coordinating the packaging of newly synthesized viral RNA into nascent immature virions.

**Figure 5:**
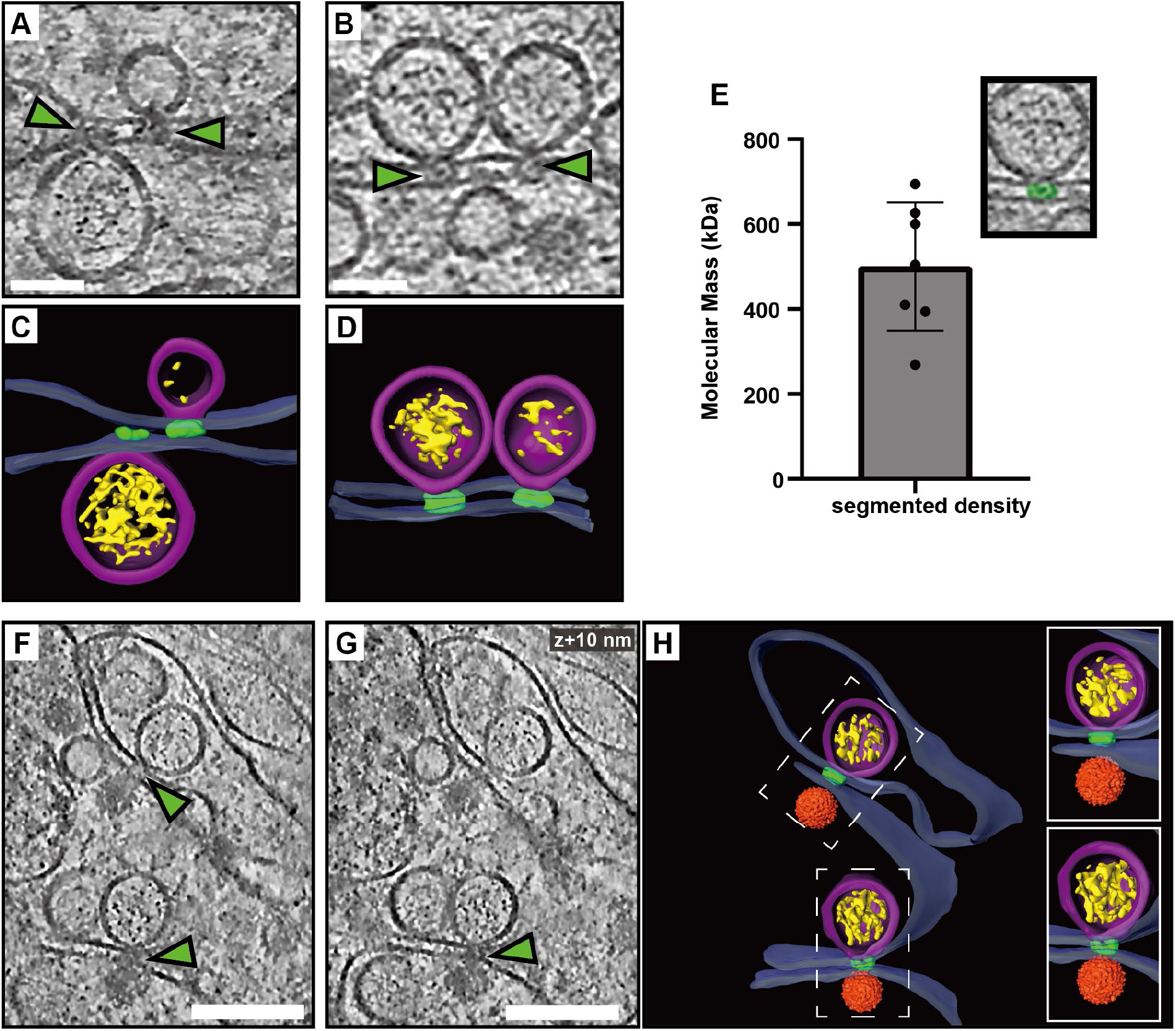
A protein complex zippers replication organelles to an apposed ER membrane. (A-B) Slices through tomograms of LGTV-infected cells showing complexes (green arrows) located at the neck of the ROs, connecting them to the adjacent ER membrane. (C-D) Segmentation of the tomograms in (A-B). (E) Estimated molecular masses of the complex (n=7). (F-G) Slices through the same tomogram at two different Z heights, showing complexes linking the RO to the site of virus assembly (green arrows). (H) Segmentation of the tomogram shown in (F-G). (C-D,H) Blue, ER membrane; purple, RO; yellow, luminal densities; green, neck complex; orange, immature virions. Scale bars, 100 nm.

### Cryo-ET reveals structural signatures of LGTV replication in *ex vivo* mouse brain

We next wished to explore if the structural features of LGTV replication that we observed in cell lines can also be identified directly in a complex, infected tissue. To do so, we proceeded to set up a workflow for cryo-ET of LGTV-infected mouse brain tissue. We based our approach on our recent publication, in which we imaged entire, *ex vivo*, LGTV-infected brains from type I interferon receptor knockout (*Ifnar*^−/−^) mice using fluorescence optical projection tomography^39^. In the 3D volumes of infected brains, immunofluorescence staining against LGTV NS5 protein was particularly strong in the choroid plexus (ChP) regions of the brain (Fig. 6A). ChP is an anatomical substructure responsible for secreting cerebrospinal fluid (CSF) into the ventricles, and as such it interfaces both with the blood and the CSF-filled ventricles (Fig. 6A). The ventricular side of the CSF-producing ependymal cells is covered with cilia and microvilli (Fig. 6A). Based on the consistently strong infection of the ChP, we developed a cryo-ET workflow for imaging ChP that was surgically removed from infected brains *post mortem*. Due to the thickness of this sample, we opted for vitrification through high-pressure freezing. The vitrified tissue was trimmed using a cryo-ultramicrotome after which lamellas were milled in place and subjected to cryo-ET (Fig. 6A). The tomograms of ChP revealed a multitude of subcellular structures such as mitochondria, nuclear pore complexes and a centriole (Fig. S6A-B). In addition, some tomograms contained virus-related features consistent with the observations in infected cell lines. This included mature virus particles encapsulated in vesicles close to membranes containing several ROs (Fig. 6B-D). Another tomogram contained a *bona fide* empty RO with seemingly thicker membrane than its limiting ER membrane, mirroring the features seen in cellular tomograms (Fig. 6E-F). Taken together, we present the first cryo-ET data on virus replication in brain tissue. The data support our cell line-based findings of RO-proximal virion maturation, and the presence of empty ROs, and provide a proof of principle that neurotropic virus replication can be studied by cryo-ET directly in infected brain samples.

**Figure 6:**
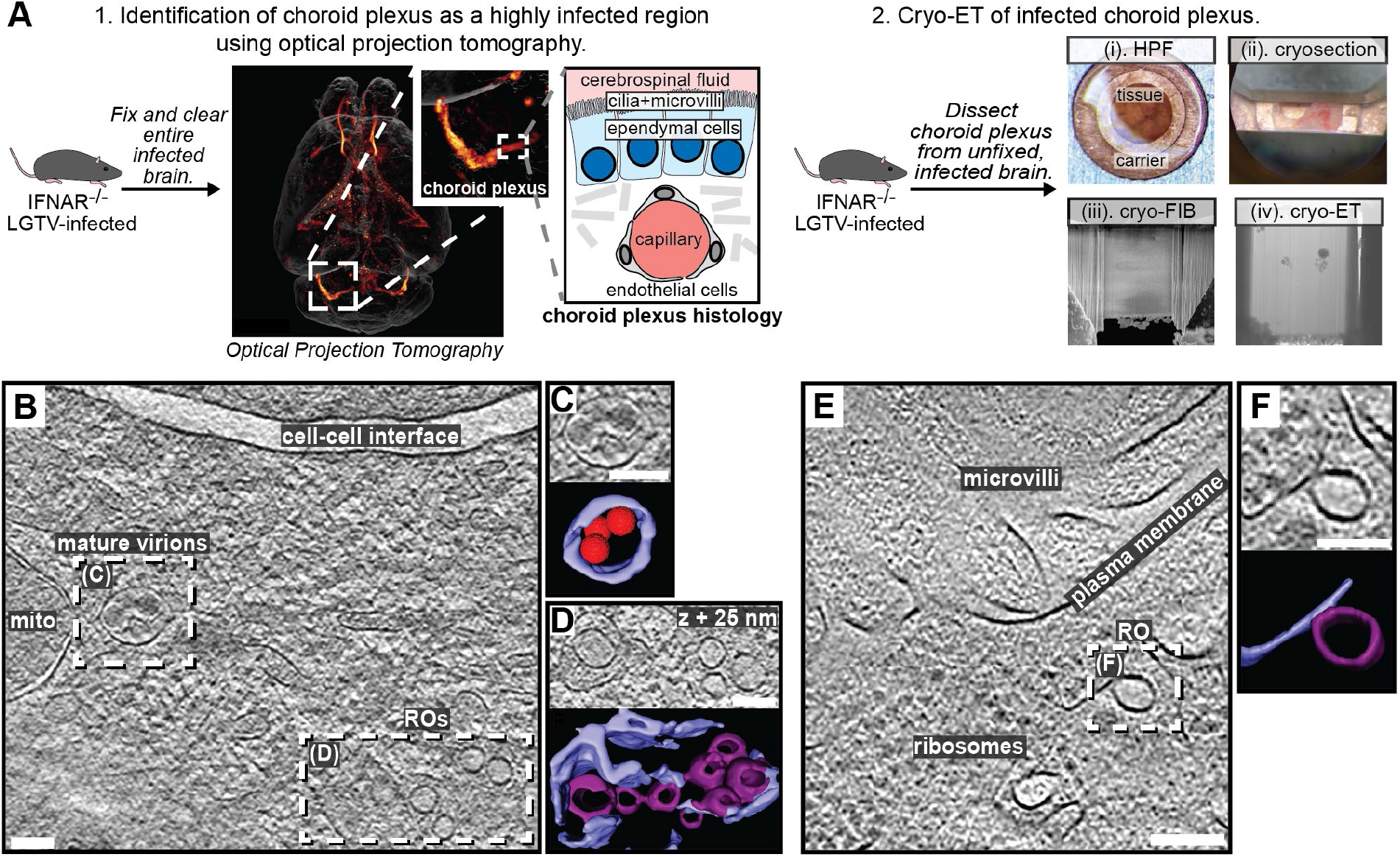
LGTV replication visualized in *ex vivo* mouse brain using cryo-ET. (A) Workflow for cryo-ET of *ex vivo* mouse brain. LGTV-infected brains from *Ifnar*^-/-^ mice were processed differently based on the imaging method. 1. Fixed and cleared brains were immunostained for the viral protein NS5 (orange/red) and imaged using optical projection tomography. Zoomed insets show infected regions of the choroid plexus, and its anatomy. 2. For cryo-ET, the choroid plexus from unfixed and unstained LGTV-infected brains was (i) high-pressure frozen, (ii) trimmed using a cryo-ultramicrotome, (iii) FIB milled, and (iv) transferred to a cryo-TEM for cryo-ET. (B) Slice through a tomogram of LGTV-infected choroid plexus showing viral replication organelles and smooth virions in proximity, enclosed within membrane vesicles. (C-D) Close-up of the areas indicated in (B) along with their corresponding segmentations. (E) Slice through a tomogram of LGTV-infected choroid plexus showing a *bona fide* RO with a thicker membrane than the ER, consistent with observations from Fig. 2. (F) Close-up of the area indicated in (E) along with its corresponding segmentation. (B-F) Colors as in Fig. 3. Scale bars, 100 nm.

## Discussion

Here, we present an *in situ* structural study of tick-borne flavivirus replication, using cryo-ET of infected cells and mouse brains. Flaviviruses are part of the vast phylum *Kitrinoviricota*, which is characterized by ROs housed in membrane buds^40^. These viruses thus need to encode mechanisms for remodeling host-cell membranes into a high-curvature bud, which is a high-energy and normally transient membrane shape. However, the viruses need to stabilize this bud-shaped membrane throughout hours of viral RNA replication. We recently showed that another genus of *Kitrinoviricota*, alphaviruses, stabilize their RO membrane through a coat-free mechanism that involves bud neck constraint by a viral protein complex, and inflation of the membrane bud by the intraluminal pressure from the viral RNA^32^. With this mechanism, the size of the replication organelle is determined by the amount of encapsulated RNA, and there are no membrane buds in the absence of intraluminal RNA. Here, we show that flaviviruses employ a different mechanism to shape the RO membrane. A membrane coat establishes a baseline RO size, allowing for the existence of ROs without intraluminal RNA. The tomograms do not indicate an ordered protein coat lattice on the RO membrane, nor clear individual protein densities, but it is possible that the curvature-generating proteins are too small and/or irregularly distributed to be detected by cryo-ET. We suggest that a strong candidate for this RO curvature generator is the 39 kDa, ER lumen-resident, non-structural protein NS1. NS1 is a peripheral membrane protein that has been reported to remodel liposomes^41^, and reshape the ER membrane when overexpressed on its own^42^. The increase in RO size due to intraluminal RNA was back-calculated to stem from the pressure of a single viral genomic RNA copy in dsRNA form (Fig. 2G, Fig. S2). Alphavirus ROs also contain a single genome copy, indicating that this may be a conserved feature across *Kitrinoviricota*^32^. Whether the smaller, empty ROs represent assembly intermediates, disassembly intermediates, or dead-end, failed RO assembly events remains to be determined.

A close coupling of flavivirus genome replication and particle budding has been suggested by several lines of evidence^14,18-20^. Our cryo-electron tomograms clearly resolved the maturation state of individual virions, as supported by the good agreement between the cellular subtomogram averages and structures of purified virions^23,25^ (Fig. 3D-E, Fig. S4). The tomograms frequently revealed immature virions in the immediate vicinity of ROs, sometimes in membrane compartments opposite to ROs (Figs. 3,6). The observation of a ∼500 kDa protein complex, connecting the membranes from which ROs and virions form (Fig. 6), shows that flavivirus ROs have a “crown” or “neck complex” akin to those identified for e.g. coronaviruses, nodaviruses and alphaviruses^30,32,43-50^. Indeed, of the seven flavivirus non-structural proteins, four are integral membrane proteins of unknown structure and no known enzymatic function. Thus, it is possible that these proteins server a structural role in organizing a membrane-connecting neck complex that coordinates replication and assembly, but larger tomographic data sets would be required to get a decisive subtomogram average from infected cells. Recent publications have highlighted the potential in small-molecule antivirals that target non-enzymatic functions of flavivirus NS proteins^51-53^. These antivirals were discovered without structural insights into their target proteins, but it is possible that the neck complex we identify here is their target. Either way, a more detailed understanding of the neck complex structure and function may aid the design of improved antiviral strategies.

Contrary to the prevailing model, we observed that furin-dependent virus maturation takes place in the immediate vicinity of ROs (Fig. 3). Thus, the entire sequence replication-assembly-maturation is more closely colocalized than previously thought. The maturation compartments are distinct but intertwined with ROs (Fig. 3), suggesting a revised model of flavivirus maturation, in which a virus-induced reorganization of the secretory pathway places Golgi-like maturation compartments in the immediate proximity of ROs. Studying naturally occurring furin site variants, we could show that a single residue in the distal cleavage site affects the RO-proximal virion maturation, while having no effect on virus release and only a minor effect on lethality in an immunocompromised mouse model (Fig. 4). This suggests that the replication and infection of TBEV is robust to variations in its maturation pathway. A further step towards bridging structural and organismal studies of flavivirus replication is taken by the workflow we present for cryo-ET of infected *ex vivo* mouse brain tissue. Tomograms of infected choroid plexus revealed clear structural signatures of ROs and clustering mature virions, similar to those in cell lines (Fig. 7). While cryo-ET has recently been used to study Alzheimer’s disease in human brain^54^, our data are, to the best of our knowledge, the first cryo-ET visualization of infection processes in the brain. Future incorporation of novel lift-out and serial milling techniques into this workflow will allow for faster acquisition of larger cryo-ET data sets on infected brains^55,56^. This may e.g. enable structural analysis of virion maturation in brain samples, and comparison of replication features between different knock-out mice. In conclusion, our study identifies several novel structural features of tick-borne flavivirus replication, and places them in a cellular context that reveals a high degree of spatial coordination of genome replication, virion assembly and virion maturation.

## Materials and Methods

### Cell line and culturing

The human A549 lung epithelial cell line was grown in DMEM medium supplemented with 10% fetal bovine serum (FBS) and Penicillin Streptomycin GlutaMAX Supplement (Gibco) at 37°C in a 5% CO_2_ environment.

### Sample preparation for cryo-electron tomography of cells

Ultrafoil Au R2/2 200 mesh grids (200 mesh, Quantifoil Micro Tools GmbH) were glow discharged. Under laminar flow the grids were dipped in ethanol before being placed in µ-Slide 8 Well Chamber (IBIDI) wells. DMEM medium with 10% FBS was added to each well and incubated while cells were being prepared. Fresh medium was added to wells and cells were seeded out at 1.5x10^4^ cells/well. The seeded cells were then placed in a 37°C, 5% CO_2_ incubator for 24 h. The medium in the wells was then replaced with serum-free DMEM and either Langat virus wildtype (LGTV), recombinant chimeric R86, or recombinant chimeric Q86 added at an MOI of 20. The cells were incubated for one hour before replacing the medium with 2% FBS in DMEM and left to incubate for 24 h. The medium was replaced with fresh DMEM including 2% FBS before being taken for freezing. Plunge freezing into an ethane/propane mix was performed with a Vitrobot (ThermoFisher Scientific) at 22°C, 100% humidity, blotting time of 5 s, and blot force of -5.

### Preparation of cryo-lamellas of cells

Lamellas were milled from plunge-frozen cells using the Scios dual beam FIB/SEM microscope (ThermoFisher Scientific). Samples were first coated with a platinum layer using the gas injection system (GIS, ThermoFisher Scientific) operated at 26°C and 12 mm working distance for 10 seconds per grid. Lamella were milled at an angle range of 16°-20°. The cells were milled stepwise using a gallium beam at 30kV with decreasing current starting at 0.5 nA for rough milling and ending at 0.03 nA for final polishing of the lamella. Lamellas were milled to a nominal 200 nm thickness and stored in liquid nitrogen for less than a week before being loaded into a Titan Krios (ThermoFisher Scientific) for data collection.

### Cryo-ET data collection on cells

Data were collected using a Titan Krios (ThermoFisher Scientific) at 300 kV in parallel illumination mode. Tilt series acquisition was done using SerialEM^57^ on a K2 Summit detector (Gatan, Pleasanton, CA) in super-resolution mode. The K2 Summit detector was fitted with a BioQuantum energy filter (Gatan, Pleasanton, CA) operated with a 20eV width silt. Areas to be imaged were selected from low-magnification overview images based on the presence of convoluted cytoplasmic membranes. Tilt series were collected using a 100 µm objective aperture and a 70 µm condenser 2 aperture, after coma-free alignment done using Sherpa (ThermoFisher Scientific). Tilt series were collected using the dose-symmetric scheme with a starting angle of -13° to account for lamella pre tilt. The parameters used for acquisition were: 33,000x nominal magnification with a corresponding object pixel size of 2.145 Å in super-resolution mode, a tilt range of typically -50° to +50°, defocus between -3 µm and -5 µm, tilt increment of 2°, and a total electron dose of 110 e/ Å^2^.

### Image Processing

Motion correction, tilt series alignment, CTF estimation and correction, and tomogram reconstruction was performed as described previously^58^, using MotionCor2^59^ with Fourier binning of 2, IMOD^60,61^, and CTFFIND4^62^. For visualization and segmentation, tomograms were 3 times binned using IMOD, resulting in a pixel size of 12.87 Å. Tomograms were denoised using cryoCARE^63^ or IsoNet^64^, occasionally combined with a non-local means filter as implemented in Amira (Thermo Fisher Scientific). Segmentation of tomograms was performed in Amira, with initial membrane tracing and segmentation done using MemBrain V2^65^. Subtomogram averages of mature and immature virus particles were incorporated into segmentation using UCSF Chimera^66^. Amira was used for counting of visually recognizable features (empty and filled ROs, immature and mature particles), measurements of the distances between them, and the volume of the neck complex densities. The neck complex volumes were converted to estimated molecular masses assuming assuming 825 Da/nm^3 67^.

### Subtomogram averaging of virions

From tomograms generated with WARP^68^ at 10Å/px object pixel size, immature and mature particles were manually picked based on their clearly distinguishable appearance. 84 immature and 51 mature particles were extracted from 11 and 3 tomograms, respectively, with a box size of 80*80*80 voxels. Subtomogram averaging was done in Dynamo^69,70^, following the same procedure for both the immature and mature data sets. Initially, all particles were translationally and rotationally aligned to a single, high-contrast particle from the respective data sets, without symmetrization. These C1 averages were manually rotated and saved in UCSF Chimera^66^ to approximately fit Dynamo’s icosahedral convention, after which they were used as a template for a Gold-standard alignment with imposed icosahedral symmetry, using Dynamo’s Adaptive bandpass Filtering function. Gold-standard Fourier shell correlation curves estimated the resolution at a cutoff of 0.143 to 33 Å for the immature particles, and 80Å for the mature particles, respectively. The final averages were filtered to this resolution and masked using the spherical alignment mask.

### Estimating RO intraluminal pressure

In recent work^32^, we demonstrated the relation between the length of an RNA and the volume of the surrounding spherule. Spherules with a volume of *V* = 10^3^ − 2 ⋅ 10^3^*nm*^3^ contain 4 ⋅ 10^3^ − 10^4^ base pairs (Fig. S2A), where the number of base pairs *N* is well described by

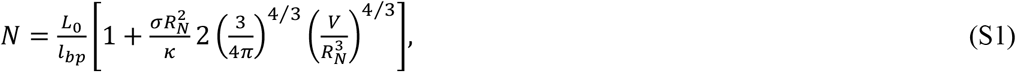

with *L*_0_ = 333*nm*, 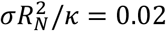, *R*_*N*_ = 9.6*nm* and the length per base pair *l*_*bp*_ = 0.256*nm*. Furthermore, a theoretical model was used to determine the relationship between the scaled pressure 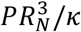 and the scaled volume 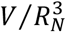 (Fig. S2B). Combining both results, we obtain a relation between the pressure *P* and the number of base pairs *N*, which shows that an RNA with a length of 2 ⋅ 10^3^ − 10^4^base pairs corresponds to a pressure of 10^−3^ − 10^−4^*k*_*B*_*Tnm*^−3^ (Fig. S2C).

### Immunofluorescence staining

Cells were grown on cover glasses and infected with LGTV as for cryo-ET. At 24 h p.i., cells were fixed with 4% formaldehyde for 20 min at room temperature and then rinsed with PBS. The fixed cells were permeabilized with 0.1% Triton X-100 in PBS for 10 min at room temperature and then rinsed with PBS. The cells were then blocked with 2% BSA in PBS containing 0.05% Tween-20 (PBS-T) for 1 hour at RT. The cells were then stained with primary antibodies against furin (goat polyclonal, dilution 1:50, Invitrogen), GM130 (mouse monoclonal clone 35/GM130, dilution 1:300, BD Transduction) or dsRNA (mouse monoclonal clone J2, dilution 1:1000, Scicons, Nordic MUbio; conjugated to allophycocyanin, Abcam) and secondary fluorescent antibodies (donkey anti-goat Alexa Fluor 488, dilution 1:1000, Invitrogen; goat anti-mouse Alexa Fluor 568, dilution 1:1000, Invitrogen) and DAPI diluted in blocking buffer for 1 hour each. Fluorescence images were acquired using a Leica SP8 confocal microscope with a HC PL APO 63x/1.4 oil CS2 objective (Leica). Confocal fluorescence images were analyzed using ImageJ Fiji software^71^.

### Chimeric viruses

A detailed description of chimeric LGTV (rLGTV^T:prME^) generation, rescue, and characterization can be found in our recent manuscript^36^. In brief, the infectious clone of LGTV, strain TP21 (kind gift of Prof. Andres Merits) was used as genetic background into which the prM and ecto-E from TBEV strain 93/783 (GenBank: MT581212.1) was inserted. Point mutation resulting in the R86Q amino acid substitution in the prM protein were introduced by overlapping PCR with primers; For: 5’ GGACGCTGTGGGAAAC<u>A</u>GGAAGGCTCACGGACA ‘3, Rev: 5’ TGTCCGTGAGCCTTCC<u>T</u>GTTTCCCACAGCGTCC 3’ (Sigma-Aldrich). RNA was generated from linearized DNA by in vitro transcription and transfected into BHK21 cells using Lipofectamine 2000 (Invitrogen). Supernatant from transfected cells was passaged once in A549 mitochondrial antiviral signaling protein (*MAVS*^*-/-*^) cells, confirmed by sequencing and used for all downstream experiments without further passaging.

### RNA isolation and qPCR

RNA was extracted from cell supernatant using Viral RNA kit (Qiagen) according to manufacturer’s instructions. The elution volume was kept constant and cDNA was subsequently synthetized from 10 µl of eluted RNA using high-capacity cDNA Reverse Transcription kit (Thermo Fisher). LGTV RNA was quantified with qPCRBIO probe mix Hi-ROX (PCR Biosystems) and primers recognizing NS3^72^ on a StepOnePlus real-time PCR system (Applied Biosystems).

### Western blot

At indicated time points, supernatant was collected and A549 cells infected with rLGTV^T:prME^ R86 or Q86 were lysed in 350 µl of lysis buffer (50 mM Tris-HCl pH7.5 + 150 mM NaCl + 0.1% Triton X-100) complemented with 1x protease inhibitor (cOmplete™ ULTRA, Roche, Basel, Switzerland) on ice for 20 min. Following lysis, cellular debris was removed by centrifugation at 14,000 g for 10 min at 4°C. Supernatant or pre-cleared cell lysate was mixed with Laemmli buffer to final concentration 1x and boiled at 95°C for 5 min. Proteins were separated by standard SDS-PAGE and transferred to an Immobilon®-P PVDF Membrane (GE Healthcare, Chicago, IL, USA). Blots were blocked overnight at 4°C in blocking buffer (PBS + 0.05 % Tween 20 + 2% Amersham ECL Prime Blocking Reagent; Cytiva), stained with primary antibodies against NS3^73^ (chicken polyclonal, diluted 1:1500), tubulin (rabbit polyclonal, diluted 1:4000, Abcam-ab6046) or M^74^ (in-house rabbit polyclonal serum, diluted 1:500) overnight at 4°C followed by secondary antibodies (goat anti-chicken Alexa-555, donkey anti-rabbit Alexa-647 (dilution 1:2500, Invitrogen, Waltham, MA, USA) for 1 h at room temperature. Blots were visualized on Amersham™ Imager 680 (GE Healthcare).

### Enzymatic assays

Synthetic peptides (Biomatik) corresponding to the P13 to P’1 residues of prM from TBEV strain Torö (Dabcyl-GRCGK<u>Q</u>EGSRTRRG-E(EDANS)) and 93/783 (Dabcyl-GRCGK<u>R</u>EGSRTRRG-E(EDANS)) or corresponding peptides with an impaired furin recognition site (Dabcyl-GRCGKREGSRTRAG-E(EDANS), Dabcyl-GRCGKQEGSRTRAG-E(EDANS)) was used as substrate. Cleavage efficiency was assayed using an adapted fluorogenic peptide assay^75^. For furin, 3 U furin (Thermo Fisher, Waltham, MA, USA) was mixed with 100 µM of substrate in a total volume of 100 µl reaction buffer (100 mM HEPES pH7.5 + 1 mM CaCl2 + 0.5% Triton X-100) at 30°C for 3h. For proprotein convertase 1/3 (PC1/3), 1 µg recombinant human PC1/3 (R&D Systems, Minneapolis, MN, USA) was mixed with 100 µM of substrate in a total volume of 100 µl reaction buffer (25 mM MES pH 6.0 + 5 mM CaCl2 + 1% (w/v) Brij-35) at 37°C for 1h. The emission at 490 nm measured every 3 min and the average rate calculated by linear regression.

### Virus infection of mice

All animal experiments were conducted at the Umeå Centre for Comparative Biology (UCCB), under approval from the regional Animal Research Ethics Committee of Northern Norrland and the Swedish Board of Agriculture, ethical permit A9-2018, A41-2019, and conducted as described previously^39,72^. Briefly, *Mavs*^-/-^ mice in C57BL/6 background (kind gift of Nelson O Gekara, Umeå University) were infected by intraperitoneal injection of 10^4^ focus-forming units (FFU) or intracranial injection of 10^2^ FFU of rLGTV^T:prME^ R86 or Q86 diluted in PBS. *Ifnar*^−/−^ mice were intracranially inoculated with 10^4^ FFU of LGTV and sacrificed when they developed either one pre-defined severe sign or at least three milder signs. Mice were monitored for symptoms of disease and euthanized as previously described criteria for humane endpoint^39^.

### Cryo-ET of LGTV-infected mouse brain tissue

Based on optical projection tomography that visualized the infection distribution in entire, *ex vivo* brains^39^, choroid plexuses were surgically removed from brains of LGTV-infected *Ifnar*^−/−^ mice *post mortem*. The choroid plexuses were perfused with ice-cold phosphate-buffered saline (PBS) and rapidly transferred to ice-cold artificial CSF^76^. Just prior to high-pressure freezing, the tissue was placed in a 3 mm copper high-pressure freezing carrier (Wohlwend) which can be clipped into an Autogrid. The sample was covered with a 20% dextran solution in PBS as cryoprotectant and covered with a sapphire disk. The assembled carrier was rapidly vitrified using a Leica EM HPM100 high-pressure freezer. The frozen carrier was trimmed at cryogenic temperatures using a Leica EM FC7 cryo-ultramicrotome with a diamond knife. The copper carrier was trimmed to leave a flat tissue sample on the carrier, measuring 100 μm in width, 20 μm in thickness, and 30 μm in depth^77^. Frozen carriers were clipped into Autogrids (ThermoFisher) prior to cryo-FIB milling with a Scios dual-beam FIB/SEM microscope (ThermoFisher Scientific).

The sample was coated with a protective platinum layer using a gas injection system for 15 seconds at a working distance of 7 mm. The cryostage was tilted at an angle of approximately 10° for milling. A rough milling was initially performed with an ion beam accelerating voltage of 30 kV and a current ranging from 0.79 to 2.5 nA to reach a thickness of 1 μm. Additionally, the two sides of the lamella were milled above and below to allow cryo-ET data collection by preventing the thick edges of the tissue from obstructing transmission EM imaging^77^. After rough milling, one edge of the lamella was detached from the main platform to relieve stress. When the lamella reached a thickness of approximately 1 μm, the ion beam current was lowered to 80 to 230 pA for fine milling, resulting in a final lamella thickness of around 200 nm. SerialEM was used to collect tilt series data with tilt angles ranging from 40° to -40° in 2° increments. The total electron dose for a single tilt series was approximately 100 e-/Å^2^, with defocus between -5 and -10 μm. Tomograms were generated as described above for cells.

### Statistics and reproducibility

Data and statistical analysis were performed using Prism (GraphPad Software Inc., USA). Details about replicates, statistical test used, exact values of n, what n represents, and dispersion and precision measures used can be found in figures and corresponding figure legends. Values of p < 0.05 were considered significant. All tomograms shown are representative of larger data sets as indicated in Table 1.

## Data availability

The cellular subtomogram averages of immature and mature Langat virus are deposited at the Electron Microscopy Data Bank with accession codes EMD-51640 and EMD-51642, respectively.

## Acknowledgements

This project was funded by a Human Frontier Science Program Career Development Award (CDA00047/2017-C to LAC), the Swedish research council (grants 2021–01145 and 2023-02664 to LAC, 2018–05851 and 2020-06224 to AÖW), a Umeå University Medical Faculty strategic grant (LAC), the Knut and Alice Wallenberg Foundation through the Wallenberg Centre for Molecular Medicine Umeå (LAC), Nadia’s Gift Foundation Innovator Award of the Damon Runyon Cancer Foundation (DRR-65-21 to DAG) and the National Institutes of Health (RF1NS125674 to DAG). SD received postdoctoral funding from the European Union under the Marie Skłodowska-Curie grant agreement No 795892. ES, NC and JZ received postdoctoral funding from the Kempe Foundation SMK-1532 (ES and JZ), and Knut and Alice Wallenberg Foundation KAW2015.0284 (NC), through the MIMS Excellence by Choice Postdoctoral Program under the patronage of Emmanuelle Charpentier. BKS received postdoctoral funding from the Wenner-Gren foundation. Cryo-EM was performed at the Umeå Center for Electron Microscopy (UCEM) a SciLifeLab National Cryo-EM facility. Fluorescence microscopy was performed at the Biochemical Imaging Center (BICU) at Umeå University. Both UCEM and BICU received funding from the National Microscopy Infrastructure, NMI (VR-RFI 2019-00217). We are thankful to all members of the VR-TBEV network, and Max Renner for valuable comments and suggestions.

## Supplementary figures and movies

**Figure S1:**
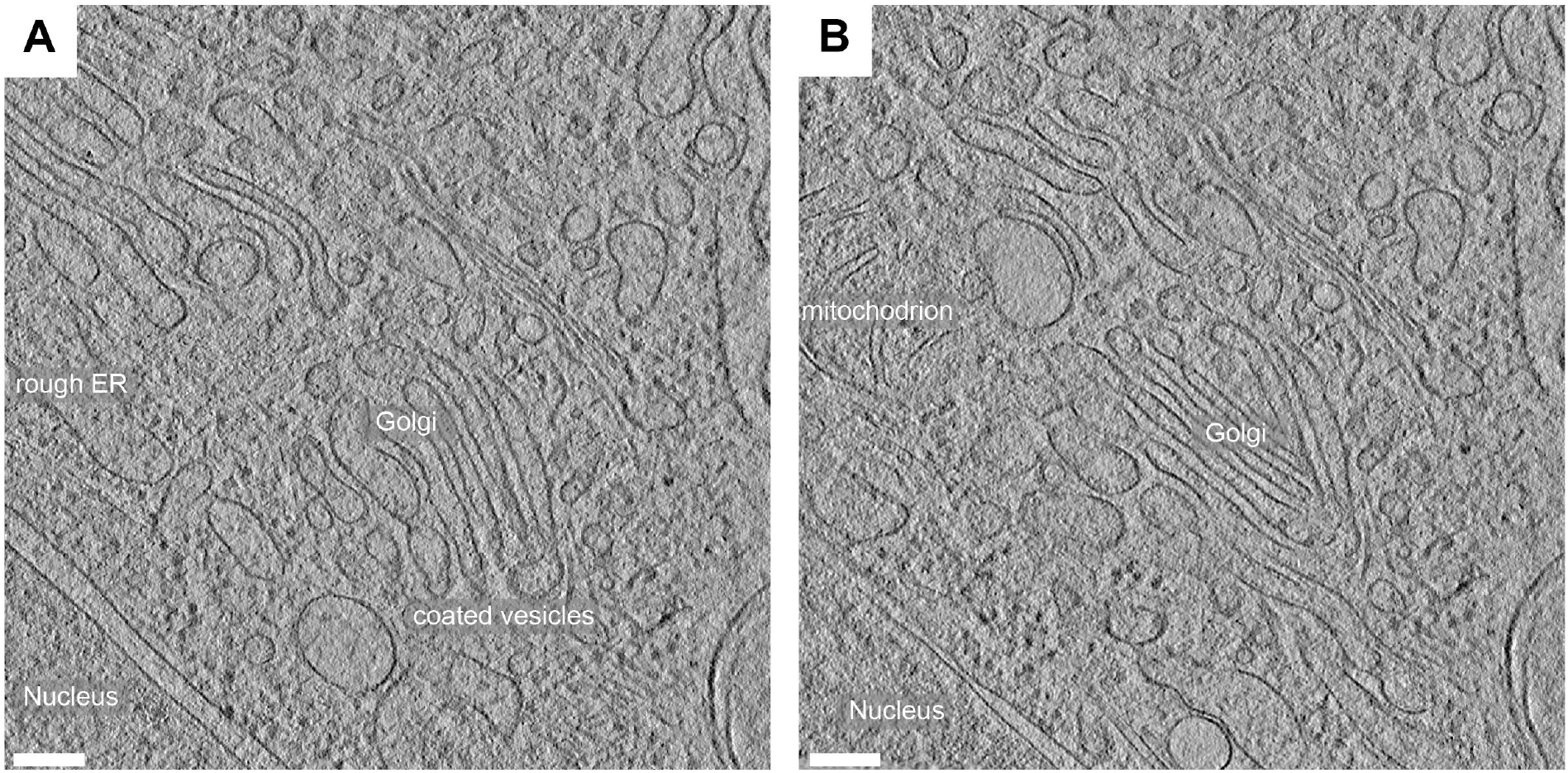
cryo-ET of uninfected A549 cells. **(A-B)** Slices from two tomograms of uninfected A549 cells reveal typical cytoplasmic features, as indicated, including a non-dilated ER and *bona fide* Golgi cisternae with typical morphology. Scale bars,100 nm.

**Figure S2.**
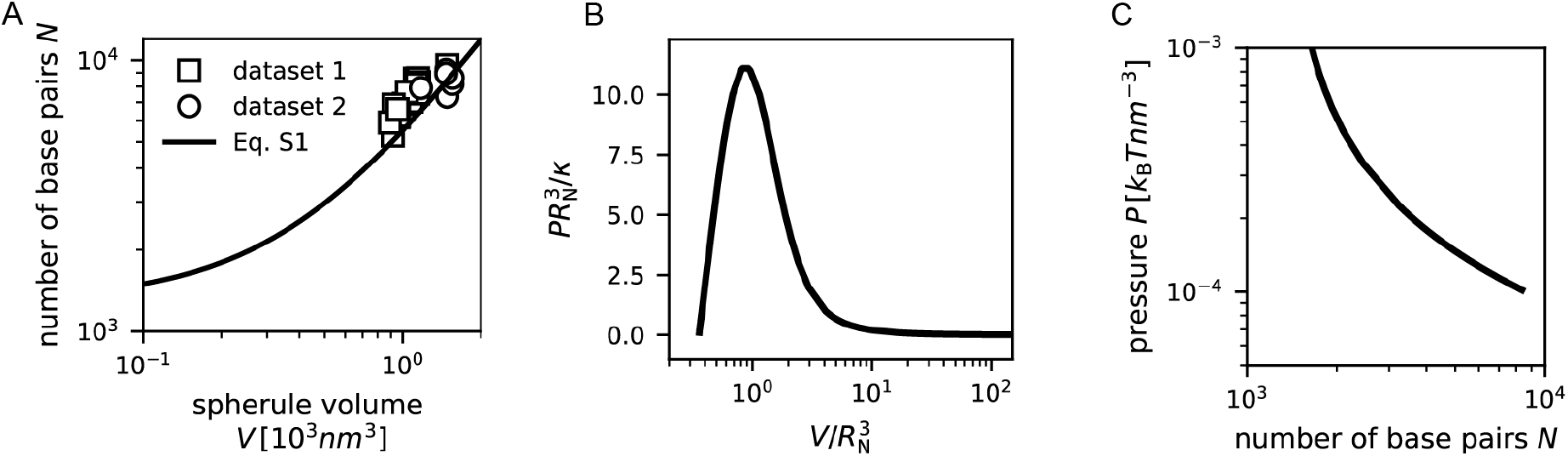
Pressure exerted by an RNA strand. (A) Relation between number of RNA base pairs and RO volume. The data is reproduced from Laurent *et al*^*32*^. We note that in Laurent *et al*^*32*^ the RNA length is shown, while here the number of base pairs is shown, assuming an interbasepair distance of 2.56 Å. (B) Relation between the scaled pressure and the scaled volume. The details of the underlying model are presented in Laurent *et al*^*32*^. (C) Relation between number of RNA base pairs N and pressure P, where we use the results from (A-B) to convert the RO volume into number of RNA base pairs.

**Figure S3:**
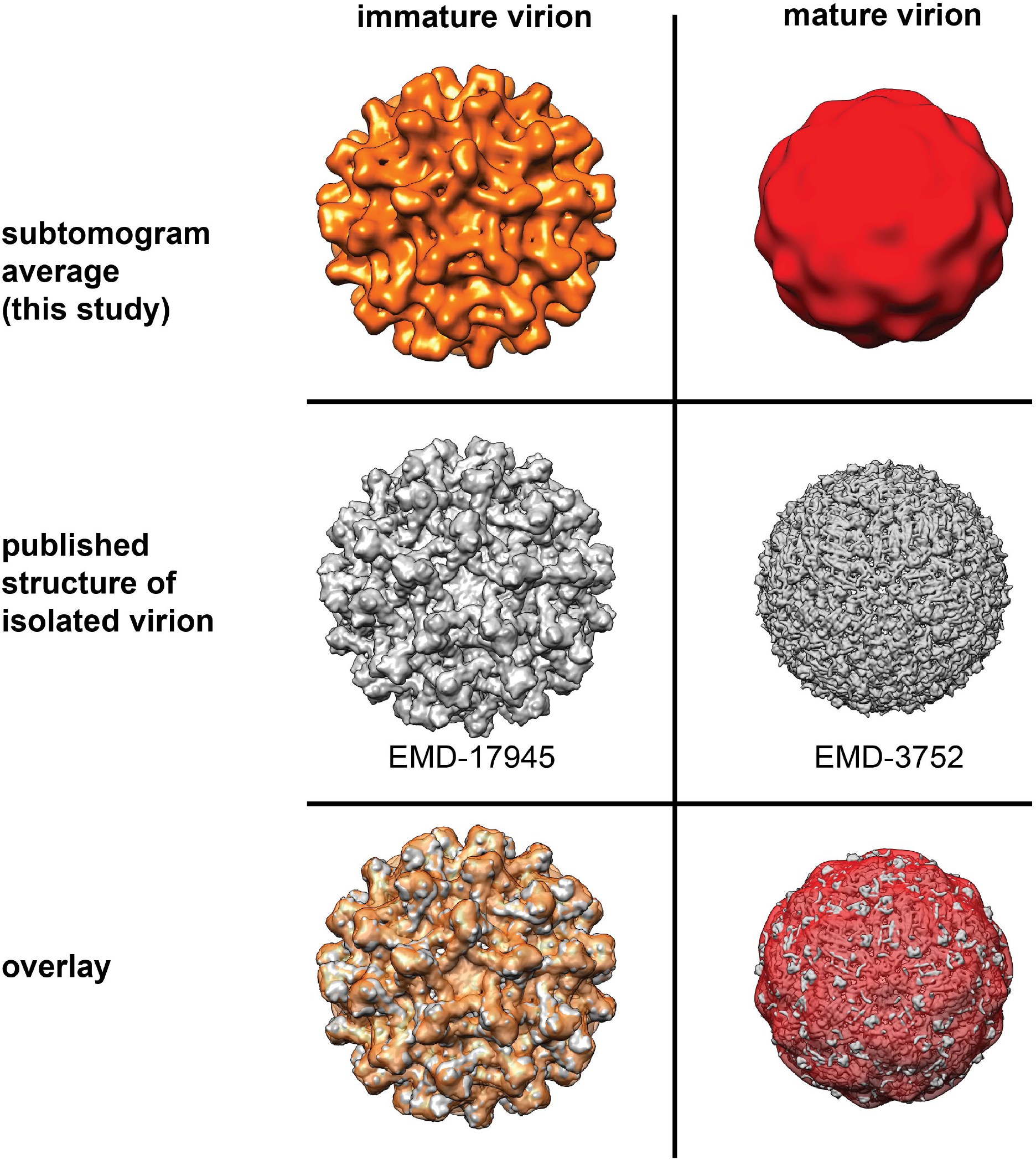
Comparison of cellular subtomogram averages with isolated virion structures. The cellular subtomogram averages from this study (top row) are compared to low-pass filtered published structures of immature and mature TBEV, from Fuzik *et al*^25^ and Anastasina *et al*^23^, respectively (mid row). The overlays (bottom row) were created using the Align to Volume command, and are shown with the subtomogram averages in semi-transparent surface representation.

**Figure S4:**
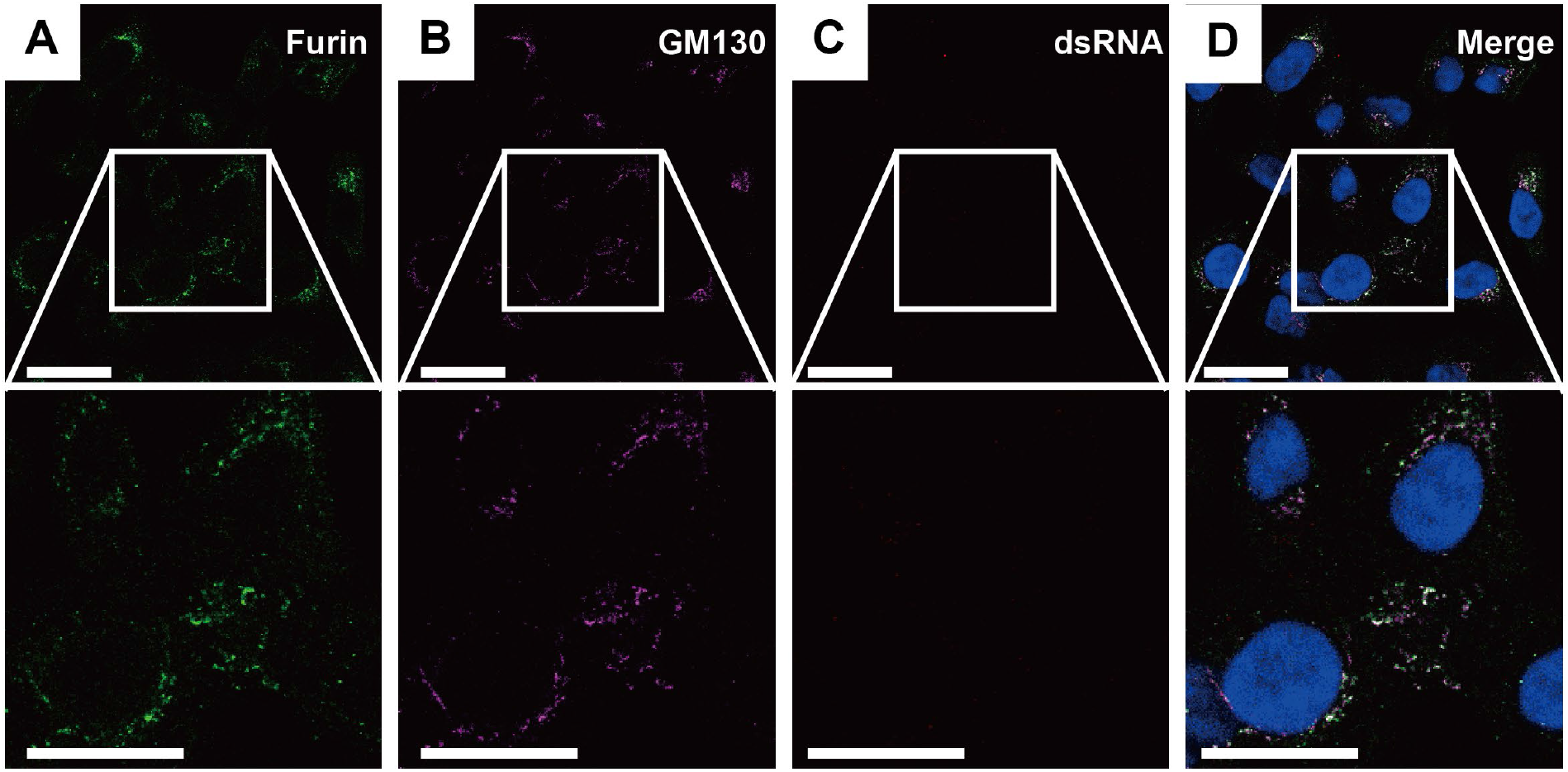
Furin localization in uninfected cells. Immunofluorescence microscopy of uninfected cells showing furin (A) and Golgi marker GM130 (B) and their colocalization in the absence of viral infection (D). Additional channels contain dsRNA staining (C) and DAPI staining of cell nuclei (D). Scale bars, 10 µm.

**Figure S5:**
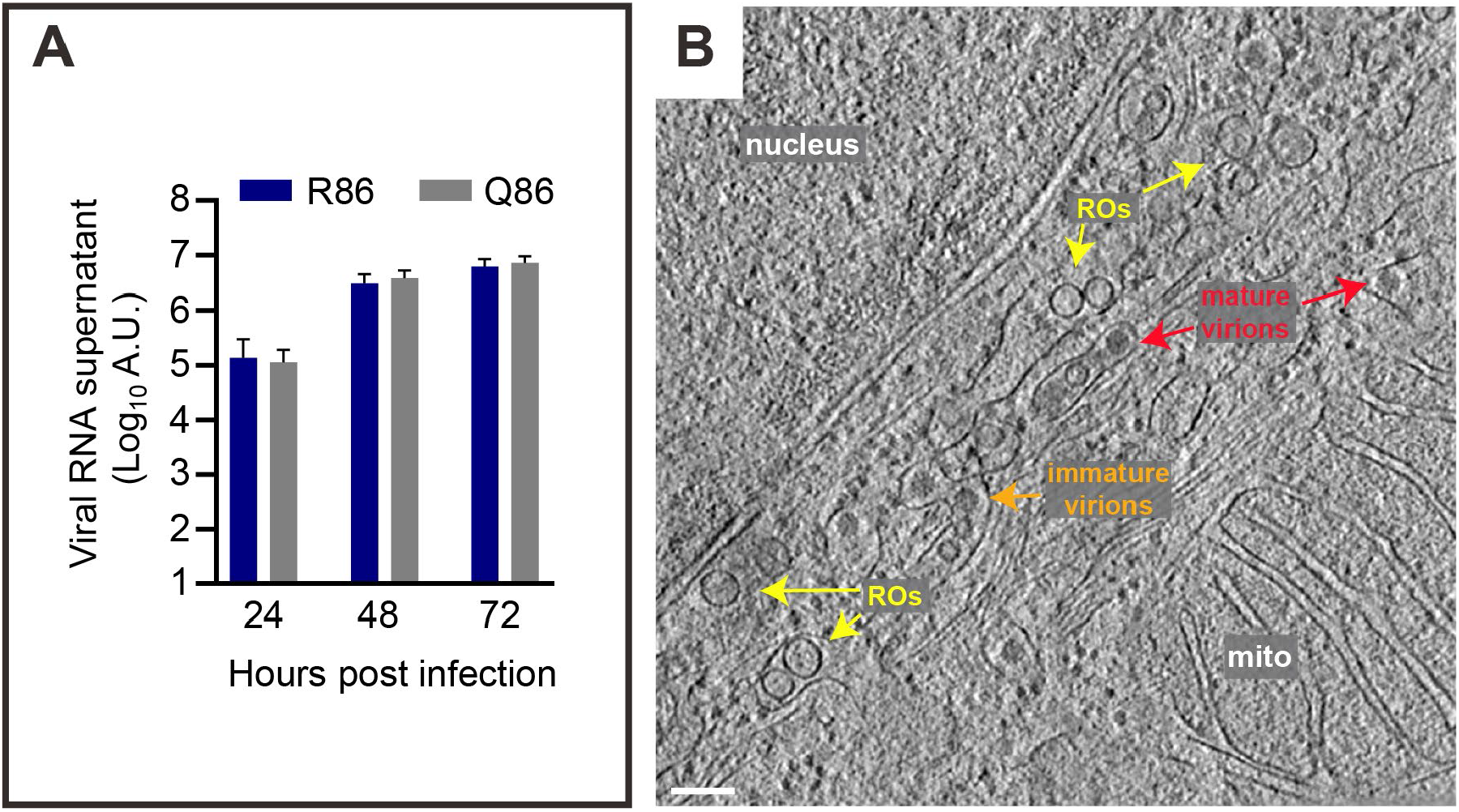
Data on rLGTV^T:prME^ Q86 and R86. (A) Growth kinetics of rLGTV^T:prME^ R86 and Q86 upon infection of A549 at MOI 1, quantitated as the amount of viral RNA in supernatant per qPCR. (B) Slice from a tomogram of a cell infected with rLGTV^T:prME^ R86 at 24 h p.i. showing various cytoplasmic and virus-related features, as indicated. Scale bar 100 nm.

**Figure S6:**
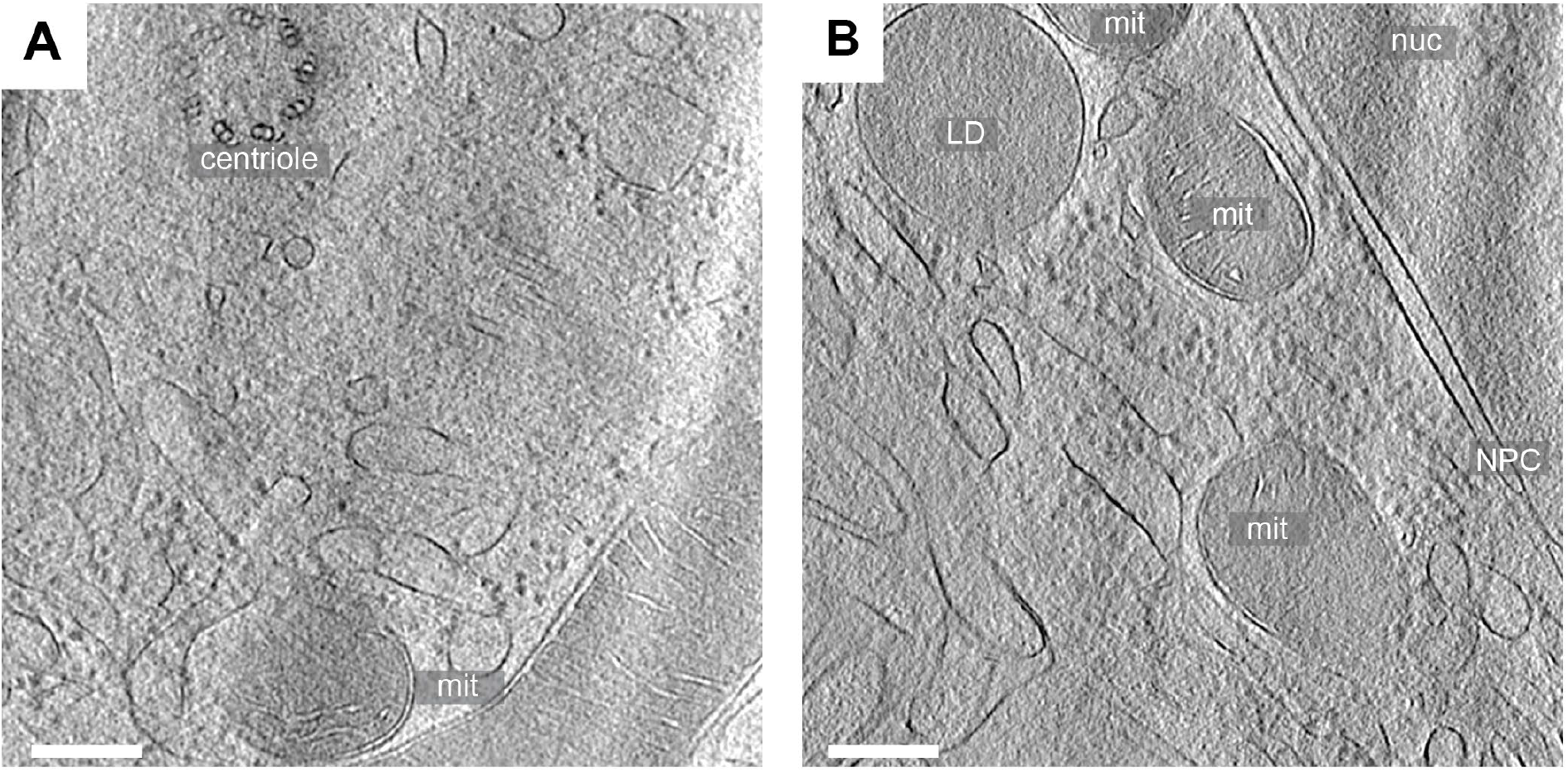
Features unrelated to infection in cryo-electron tomograms of *ex vivo* brain tissue. (A-B) Slices from two tomograms of high-pressure frozen choroid plexus from LGTV-infected *Ifnar*^-/-^ mice. The features indicated are a centriole, several mitochondria (mit), a lipid droplet (LD), the peripheral area of a nucleus (nuc) and the nuclear envelope including one nuclear pore complex (NPC). Scale bar, 100 nm.

